# Atomic elementary flux modes explain the steady state flow of metabolites in large-scale flux networks

**DOI:** 10.1101/2024.11.13.623484

**Authors:** Justin G. Chitpin, Theodore J. Perkins

## Abstract

Steady state fluxes are a measure of cellular activity under metabolic homoeostasis, but understanding how individual substrates are metabolized remains a challenge in large-scale networks. Pathway-based approaches such as elementary flux mode (EFM) analysis are limited to small networks due to the combinatorial explosion of pathways and the ambiguity of decomposing fluxes onto EFMs. Here, we present an alternative approach to explain metabolic fluxes in terms of the steady state flow of their atomic constituents. We refer to these pathways as atomic elementary flux modes (AEFMs) and show that computations involving AEFMs are orders of magnitude faster than standard EFMs. Using our approach, we enumerate carbon and nitrogen AEFMs in five genome-scale metabolic models and compute the AEFM decomposition of fluxes estimated in a HepG2 liver cancer cell line. Our results systematically characterize carbon and nitrogen remodelling and, on the HepG2 network, predict glutamine metabolism through a recently discovered non-canonical TCA cycle.

## Introduction

Recent technological and computational advancements are leading to a rise in metabolic flux data (Lee et al, 2019; Wang et al, 2022a,b). These fluxes quantify the rate of metabolite interconversions within a cell or their movements across subcellular compartments. Metabolic fluxes are studied to understand the dynamics of substrate switching (Zampieri et al, 2019; Kochanowski et al, 2021), redox balancing (McKinlay & Harwood, 2010; Franchina et al, 2022), overflow metabolism (Basan et al, 2015; Vander Heiden et al, 2009), cell differentiation (Endo et al, 2022; Jackson & Finley, 2024), and cellular communities (Schuetz et al, 2012; Yu et al, 2022; Gustafsson et al, 2024). Hence, ‘fluxomics’ (Emwas et al, 2022; Sanford et al, 2002) is emerging as a field dedicated to quantifying the totality of fluxes within biological systems.

Current experimental and computational methods generally estimate fluxes under metabolic homeostasis. Under this assumption, the production and consumption of metabolites are balanced over time, and the fluxes are said to be under steady state. Techniques such stable isotope tracing analysis (SITA) experimentally determine steady state fluxes by culturing cells in labelled substrates and quantifying their incorporation within endogenous metabolites (Buescher et al, 2015; Antoniewicz, 2018). Using this method, fluxes have been measured in major metabolic subsystems such as glycolysis, pentose phosphate pathway, and the TCA cycle (Orth et al, 2010b; Ahn et al, 2017; Wu et al, 2019). Many computational methods, including but not limited to flux balance analysis (FBA), have also been proposed to infer steady state fluxes from transcript, protein, and metabolite abundance data (Uematsu et al, 2022; González-Arrué et al, 2023; Huang et al, 2023; Kaste & Shachar-Hill, 2023; Lee et al, 2023). Collectively, these approaches are being used to estimate and predict fluxes in genomescale metabolic models (GEMs) involving hundreds to thousands of reactions (Nilsson et al, 2020b; Cherkaoui et al, 2022; Gelbach et al, 2024).

Although much effort has been made to estimate steady state fluxes, less emphasis has been placed on developing methods to analyze these large-scale flux datasets. In contrast to abundance data (e.g. transcripts, proteins, metabolites), steady state fluxes are constrained by the reaction stoichiometries between metabolite inflows and outflows. In a network with fixed source and sink fluxes, an increase in flux along one metabolic pathway must decrease fluxes along one or more other pathways to maintain mass conservation. For simple networks, one may predict how the system would evolve by identifying the up and downstream reactions flanking an enzymatic perturbation. For example, Schwartz & Kanehisa were able to enumerate all glycolytic pathway fluxes in a 12-reaction model of yeast glycolysis (Schwartz & Kanehisa, 2006). Yet in more complex genome-scale metabolic networks containing thousands of fluxes (e.g. Brunk et al (2018); Robinson et al (2020); Wang et al (2021)), there exists no hierarchical relationship between reactions. A change in one reaction rate, let alone multiple perturbations, may alter network fluxes beyond neighbouring reactions (Nobile et al, 2021).

This problem of understanding the flow of metabolites within steady state flux networks has historically been addressed through the concept of elementary flux modes (EFMs). EFMs are a pathway-based approach to analyze metabolic networks and are defined as minimal sets of biochemical reactions that maintain steady state flux (Schuster & Hilgetag, 1994). This minimal property specifies that an EFM cannot be decomposed into two or more smaller pathways carrying steady state flux. An attractive property of EFMs is that any set of steady state fluxes in a network can be explained as a positive, linear combination of EFMs (Schuster et al, 1999). While EFMs are a mechanistic explanation of how metabolites travel through networks, the number of EFMs scales combinatorially as a function of network size (Acuña et al, 2009). It remains computationally infeasible to enumerate EFMs in GEMs after over a decade of algorithmic advancements (e.g. Gagneur & Klamt 2004; Terzer & Stelling 2008; Buchner & Zanghellini 2021). The number of EFMs also poses a challenge for explaining network fluxes in terms of their EFMs. When there are more EFMs than linearly independent fluxes, there exist multiple ways to assign EFM weights that reconstruct the observed network fluxes. Consequently, structural analyses of EFMs and downstream analyses of their weights have been limited to small-scale metabolic subnetworks (Fakih et al, 2023; Mpabanzi et al, 2022; Toya & Shimizu, 2022; Matsuda et al, 2011; Schwartz & Kanehisa, 2006).

In our previous work, we developed a novel solution to the EFM flux decomposition problem for the special case of unimolecular reaction networks under a Markov constraint. We modelled a hypothetical particle moving between metabolites under the assumption that the transition probability of each reaction was proportional to the observed steady state flux. By expanding the notion of Markov chain state to include sequences of metabolites, we proposed an algorithm to efficiently compute simple cycle probabilities corresponding to a given EFM. We called this construct the *cycle-history Markov chain* (CHMC) and proved that these EFM probabilities could be scaled to weights that fully reconstructed the flux along each reaction (Chitpin & Perkins, 2023).

In this work, we generalize our CHMC method to operate on any type of metabolic network, including those involving multispecies reactions. We do so by proposing a new type of *atomic elementary flux mode* which traces the steady state flow of an individual atom within a metabolic network. We refer to these pathways as AEFMs to distinguish them from the (molecular) EFMs proposed by Schuster & Hilgetag. While a linear combination of molecular EFMs will reconstruct the overall reaction fluxes, the totality of AEFM weights for a given source metabolite will always reconstruct its mass flow entering the network. We first introduce our framework for enumerating and uniquely identifying AEFM weights. Using a state-of-the-art atom mapping algorithm, we show how to generate atomic transition graphs from which we construct atomic cycle-history Markov chains (ACHMCs). From these ACHMCs, we enumerate AEFMs in GEMs considered infeasible by state-of-the-art molecular EFM enumeration programs. We further analyse the structure of the AEFMs to characterize nutrient source remodelling. Lastly, we analyze a single set of AEFM weights estimated from a HepG2 liver cancer cell line to quantify glutamine metabolism. Our results predict glutamine in HepG2 cells is metabolized through a non-canonical TCA pathway recently discovered in non-small cell lung cancer, mouse embryonic, and C2C12 mouse myoblast cells (Arnold et al, 2022).

## Results

### Tracing atomic flows in genome-scale models

We first describe what molecular EFMs look like in metabolic networks and provide intuition why enumerating and decomposing fluxes onto these pathways remains a fundamental problem. Figure 1a shows an example molecular EFM involving substrates glucose and ATP which are consumed to produce metabolites G3P, DHAP, and ADP. Multiple source and sink fluxes are required to balance the internal metabolite stoichiometries, leading to several branching and converging paths within the molecular EFM. As the metabolic networks expands to include dozens or more reactions, the corresponding molecular EFMs can grow increasingly complex with numerous source and sink fluxes required to stoichiometrically balance participating metabolites. In particular, there may be multiple sets of reactions involved to balance pairs of energetic molecules (e.g. ATP/ADP), redox equivalents (e.g. NAD^+^/NADH), metabolic byproducts (e.g. water, hydrogen ions), or coenzymes (e.g. coenzyme A). This complexity can lead to an intractable number of pathways that may only differ by reaction subsets required to mass-balance these metabolites. Choosing certain metabolites to remove may, in part, solve this problem. However, the choice of metabolites to prune from the network is often subjective and not guaranteed to sufficiently reduce the total number of molecular EFMs.

**Figure 1:**
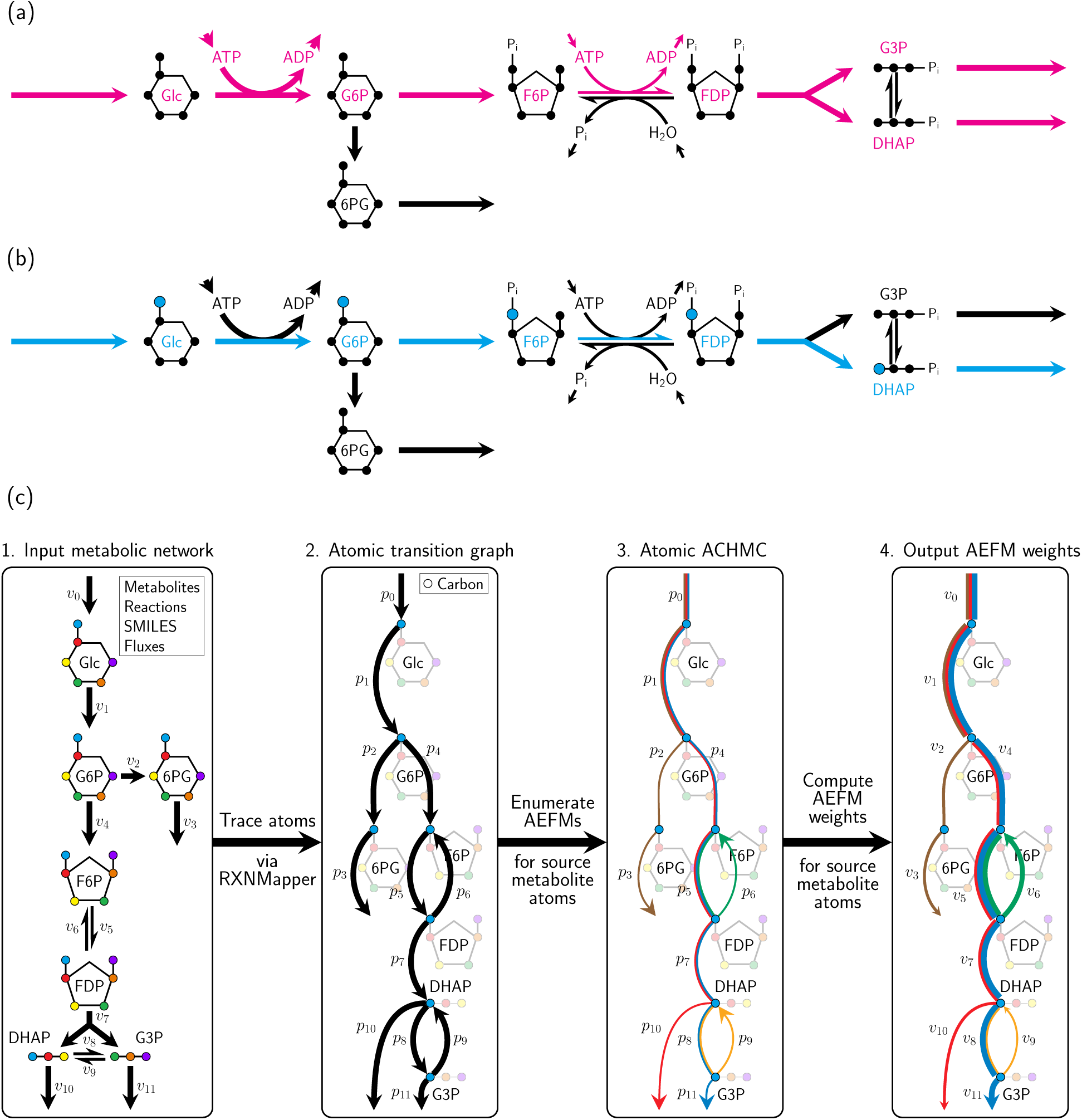
Atomic cycle-history Markov chain (ACHMC) pipeline. (a) An example molecular EFM in glycolysis. (b) An example AEFM tracing the flow of a given glucose carbon in glycolysis. (c) The major steps within our pipeline to enumerate AEFMs and compute their weights.

Given many metabolic flux networks are generated to study nutrient catabolism (Zamboni et al, 2015), we argue for studying source metabolite-derived AEFMs to reduce the combinatorial number of molecular EFMs. We highlight an example carbon AEFM in Figure 1b which shows how the C6 glucose carbon is remodelled during the first steps of glycolysis. As atoms are never split apart or fused together in metabolic reactions, these AEFMs always correspond to simple paths spanning source and sink metabolites or simple cycles between internal metabolites. These pathways are straightforward to interpret because each atom within an AEFM can only transit from a single substrate to product, regardless of the network size, connectivity, or number of participating substrates/products in each reaction. This property further enables us to enumerate and compute the AEFM weights using our previously described CHMC method to uniquely decompose steady state fluxes onto EFMs for unimolecular reaction networks (Chitpin & Perkins, 2023).

We outline our computational pipeline in Figure 1c to enumerate AEFMs and uniquely assign their weights when the steady state network fluxes are known. In step 1, we require as inputs the metabolic network defined by a set of metabolites and reactions encoded as a stoichiometry matrix, the metabolite structures denoted as SMILES strings, and the steady state fluxes. From the stoichiometry matrix and SMILES strings, we construct reaction SMILES strings which are imputed into the atom mapping program RXNMapper (Schwaller et al, 2021) to map the position of each atom within each reaction equation. This atom mapping data is used to construct an atomic transition graph in step 2, which traces the atomic position of an individual atom from a given source metabolite entering the network. We set the transition probabilities proportional to the atomic flux through each reaction, accounting for those with multiple substrate stoichiometries (see Methods and Figure S1). For each atomic transition graph, we construct the corresponding ACHMC to enumerate all AEFMs involving that source metabolite atom in step 3. Finally, the AEFM weights are computed in step 4 by steady state analysis of the ACHMC.

### Enumerating AEFMs is computationally tractable in large-scale networks

Given the major computational challenge of enumerating molecular EFMs in networks with *>* 100 metabolites and reactions, we first sought to test whether it was feasible to enumerate AEFMs in GEMs. We selected five genome-scale metabolic models across distinct organisms in ascending order of network size. These networks include a metabolic model of *E. coli* (E. coli core; Orth et al 2010b; Data ref: Orth et al 2010a), human red blood cells (iAB RBC 283; Bordbar et al 2011b; Data ref: Bordbar et al 2011a), *H. pylori* (iIT341; Thiele et al 2005b; Data ref: Thiele et al 2005a), *S. aureus* (iSB619; Becker & Palsson 2005b; Data ref: Becker & Palsson 2005a), and human liver cancer cells (HepG2; Nilsson et al 2020b; Data ref: Nilsson et al 2020a). These models were pre-processed to ensure unambiguous atom mappings across all reactions. Namely, all metabolites with no known structures (pseudometabolites), and pseudoreactions with non-integer stoichiometries were removed from the networks. The resulting metabolic networks ranged between 71-547 metabolites and 144-974 reactions (Figure S2). We then attempted to enumerate the molecular EFMs using FluxModeCalculator, which is the fastest molecular EFM enumeration program (van Klinken & Willems van Dijk, 2016). Following our pipeline, we subsequently compiled the metabolite SMILES strings and constructed ACHMCs rooted on each carbon and nitrogen atom across all source metabolites. The resulting summary statistics describing the GEMs and ACHMCs are presented in Table 1. Across all carbon ACHMCs, we observed a broad increase in the total number of AEFMs as a function of metabolic network size. The one exception to this observation was the HepG2 network which contained fewer metabolites and reactions than iSB619, yet exhibited two orders of magnitude more carbon AEFMs. Inspection of the HepG2 network revealed it only contained 40 source metabolites compared to the 198 source metabolites in the iSB619 network, indicating a greater degree of network connectivity between internal metabolites (Figure S2).

**Table 1:** Summary statistics for the five metabolic networks in this study.

| GEM (# mets \| # rxns) | Carbon metabolism |  |  |  |  |
| --- | --- | --- | --- | --- | --- |
|  | Sources | Sinks | States | Transitions | AEFMs |
| E. coli core (71 \| 144) | 184 | 255 | 14659 | 24355 | 6519 |
| iAB_RBC_283 (274 \| 465) | 954 | 1166 | 37122 | 51220 | 7958 |
| iIT341 (433 \| 759) | 1486 | 2134 | 86342 | 120545 | 30633 |
| iSB619 (547 \| 974) | 2188 | 2986 | 169877 | 246712 | 59444 |
| HepG2 (435 \| 545) | 376 | 939 | 9092000 | 11615891 | 1377937 |
| GEM (# mets \| # rxns) | Nitrogen metabolism |  |  |  |  |
|  | Sources | Sinks | States | Transitions | AEFMs |
| E. coli core (71 \| 144) | 30 | 40 | 246 | 448 | 204 |
| iAB_RBC_283 (274 \| 465) | 161 | 214 | 6409 | 9275 | 2091 |
| iIT341 (433 \| 759) | 314 | 447 | 27440 | 39761 | 10440 |
| iSB619 (547 \| 974) | 353 | 516 | 36661 | 54165 | 14214 |
| HepG2 (435 \| 545) | 72 | 190 | 19137 | 27860 | 5105 |

Figure 2a shows the total number of atomic and molecular EFMs enumerated by our program MarkovWeightedEFMs.jl and FluxModeCalculator on the five GEMs. Using our ACHMC pipeline, we enumerated between 10^3^ *−* 10^6^ carbon and 10^2^ *−* 10^3^ nitrogen AEFMs within each GEM. In contrast, FluxModeCalculator failed to enumerate molecular EFMs in all but the two smallest E. coli core and iAB RBC 283 datasets, returning 10^3^ *−* 10^4^ times more molecular-versus-atomic EFMs.

**Figure 2:**
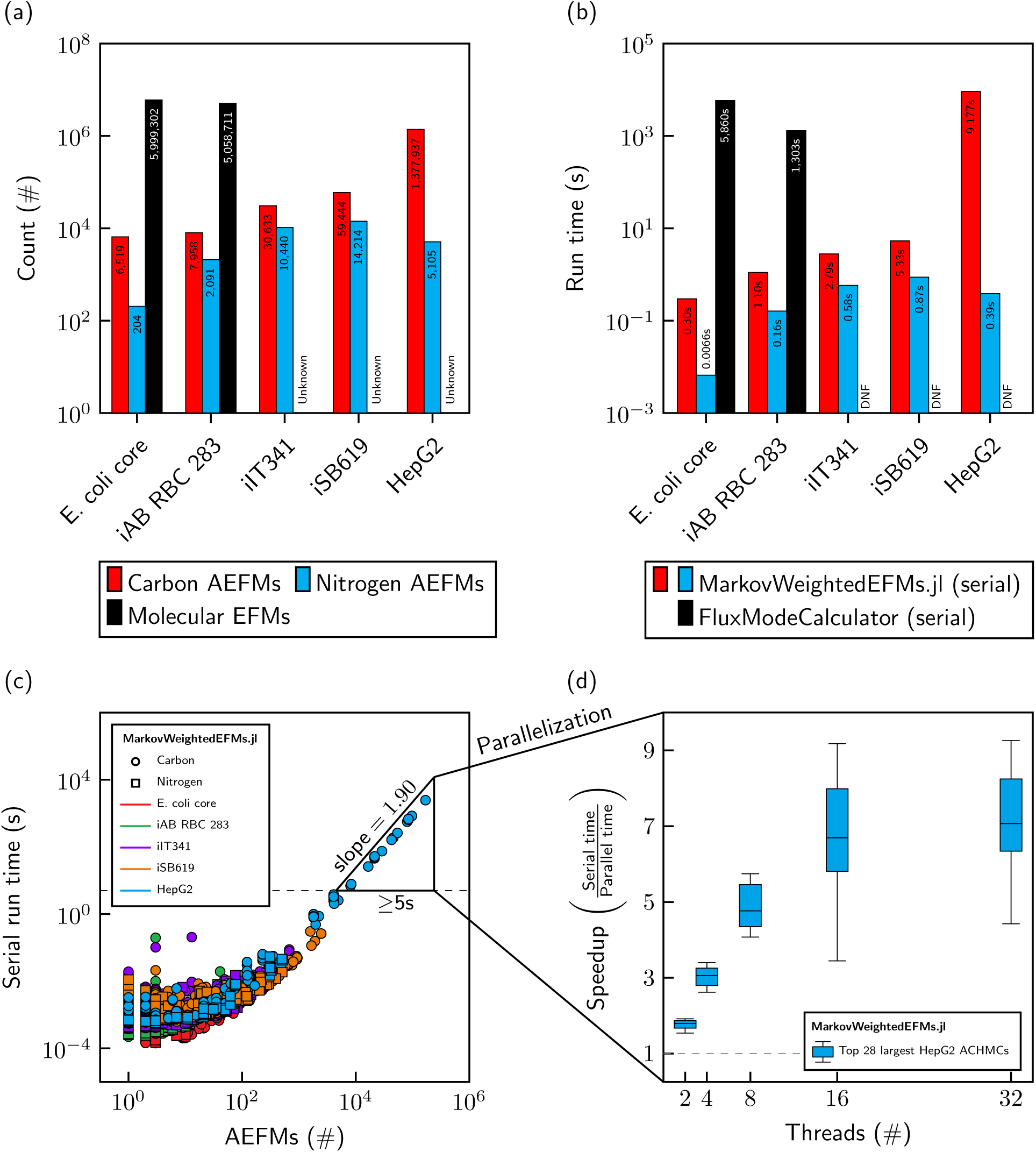
AEFM enumeration is computationally tractable compared to molecular EFMs in five GEMs. (a) Number of atomic and molecular EFMs in the five GEMs. (b) The run time of our MarkovWeightedEFMs.jl method versus FluxModeCalculator running in serial. (c) Serial run time scaling of MarkovWeightedEFMs.jl as a function of the number of AEFMs. (d) Speedup of selected ACHMCs as a function of additional threads.

By focusing on an atom of interest, our ACHMC approach greatly reduced the number of EFMs to biologically relevant pathways involving nutrient source remodelling. This reduction in pathways resulted in AEFM-versus-EFM enumeration completing 10^3^ *−* 10^4^ times faster (Figure 2b). While running FluxModeCalculator in parallel did improve run times by 8.2-8.6 times (Figure S3), it remained over 10^1^ *−* 10^2^ slower than enumerating their corresponding carbon and nitrogen AEFMs. We further found that AEFM enumeration actually involved fewer metabolite-atom combinations than the total number of metabolites in a given network. Constrained by the atom mapping predictions, we observed that the average carbon and nitrogen ACHMC across all five GEMs only transited 0.41% and 1.7% of the total number of metabolite-carbon and metabolite-nitrogen positions, respectively (Figure S4). While serial run times in Figure 2c scaled quadratically (Figure 2c; slope = 1.90, intercept = *−*6.54), our program and AEFM approach benefits greatly from parallelism. Each ACHMC can be computed independently of one another and our program is also parallized on the level of individual ACHMCs to improve AEFM enumeration. For the top 28 largest carbon ACHMCs in the HepG2 network, Figure 2d shows a median speedup of 7.1 times when using 32 threads.

### Most AEFMs are source-to-sink pathways spanning dozens of reactions

We next turned to characterizing the structural properties of all carbon and nitrogen AEFM across the five GEMs. Molecular EFMs can be divided into pathway that span source and sink metabolites or internal loops within the metabolic network such as pairs of reversible reactions. Both pathways have clear biological interpretations with source-to-sink pathways explaining the steady state flow of source metabolites on their way to exiting the network, and internally looped pathways corresponding to substrate cycles. At an atomic level, these pathways can be subclassified further into AEFMs that transit sets of distinct metabolites or those that revisit the same metabolite, albeit in different atomic positions. This notion of a “metabolite revisitation” is similar to an isotopomer in stable isotope tracing (Antoniewicz, 2018). An example source-to-sink AEFM with this property is a glucose-derived carbon AEFM remodelling through two turns of the TCA cycle; on the second turn, the position of the carbon atom shifts within the TCA intermediates, resulting in different isotopomers (Duan et al, 2022) and therefore a source-to-sink pathway with a metabolite revisitation under our nomenclature.

To obtain a broad understanding of the types of AEFMs within the five GEMs, we categorized them into source-to-sink pathways and looped pathways with or without metabolite revisitations in different atomic positions. Figure 3a-b shows the counts for each AEFM classes within each GEM. As expected, we observed that the majority of carbon AEFMs were source-to-sink pathways with or without metabolite revisitations. Those without metabolite revisitations tended to scale as a function of network size, while those with revisitations increased modestly. Investigation of these source-to-sink pathways confirmed that many metabolite revisitations involved TCA cycle intermediates. Looped AEFMs without metabolite revisitations primarily corresponded to transport reactions cycling metabolites across compartments and cyclic pathways involving TCA intermediates. These two observations explain, in part, why the iAB RBC 283 network contained the fewest number of pathways with metabolite revisitations, since they lack organelles and cannot support cyclic AEFMs associated with metabolite transport between compartments nor mitochondrial TCA activity. There were very few internally looped AEFMs with metabolite revisitations with no obvious trend across the five GEMs. The greatest number of these pathways were found in the E. coli core network with many metabolite revisitations involving carbon dioxide derived from isocitrate and alpha-ketoglutarate re-entering the TCA cycle via oxaloacetate. As there are fewer metabolites containing nitrogen, we found the majority of their AEFMs corresponded to pathways without metabolite revisitations, primarily involving amino acid remodelling or transports between compartments. Looped pathways with metabolite revisitations were only observed in the iIT341 and iSB619 networks through reactions involving the production and consumption of ammonia.

**Figure 3:**
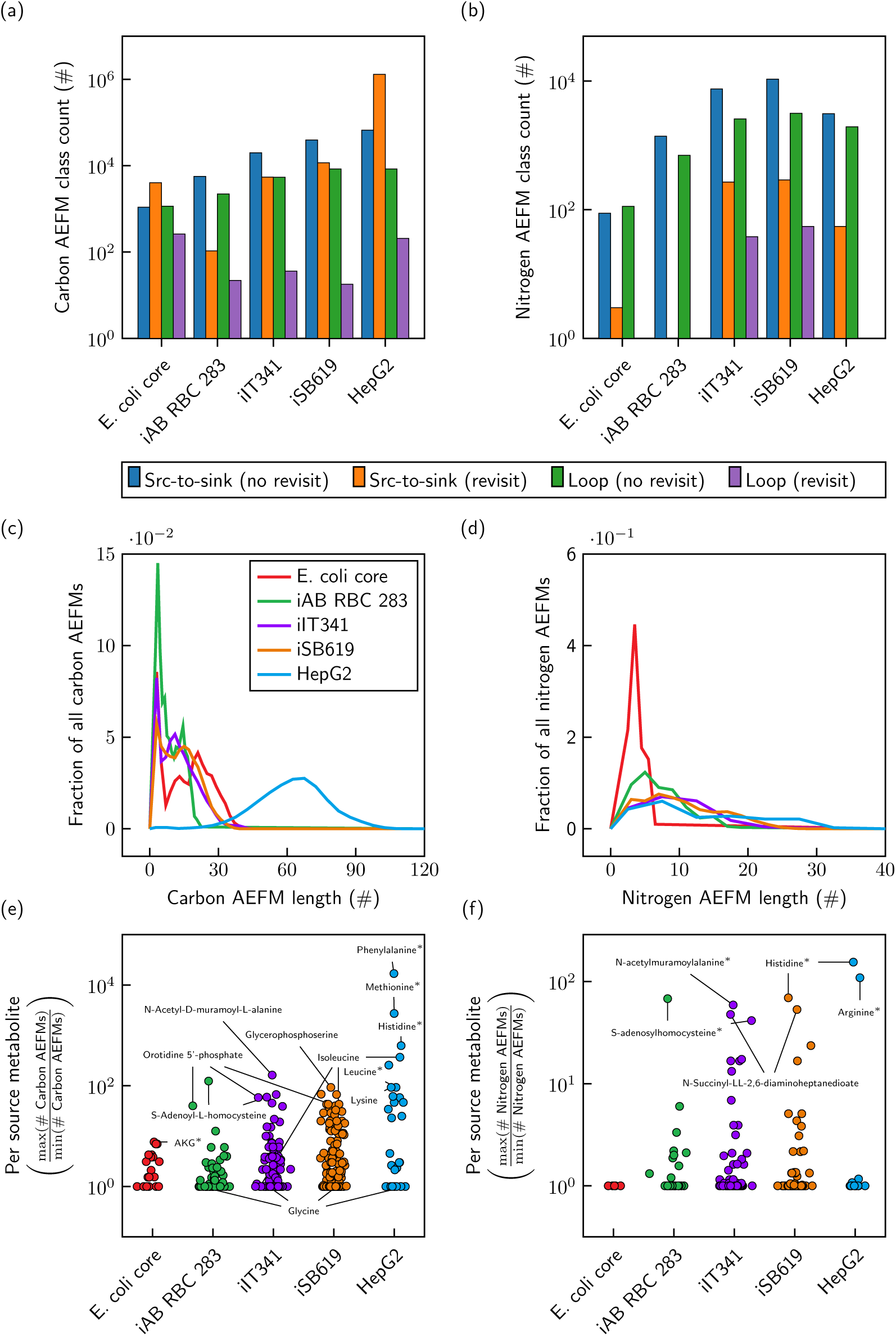
Structural analysis of AEFMs in five genome-scale metabolic models. (a-b) Classes of carbon and nitrogen EFMs. (c-d) Fraction of carbon and nitrogen AEFM lengths. (e-f) Ratio of the greatest and smallest number of AEFMs between carbons and nitrogens within the same source metabolite. Representative metabolites from each dataset are labelled; labels with asterisks indicate that metabolite is also present in other networks.

Since many looped AEFMs involve reversible reactions or metabolite transport across compartments, we hypothesized that looped AEFMs should involve fewer reactions than source-to-sink pathways which can span numerous internal metabolites. We calculated the fraction of carbon and nitrogen AEFMs with given lengths across all five GEMs and plotted their distributions in Figure 3c-d. Our results show that carbon AEFMs in the first four networks are bimodally distributed with the majority of pathways involving a dozen or fewer reactions. While carbon AEFMs of the HepG2 network appeared to exhibit a single Gaussian distribution (*µ* = 61.7, *σ*^2^ = 15.5), closer inspection of the histogram revealed a small peak spanning 2-12 reactions. To determine whether these peaks were associated with source-to-sink and looped AEFM classes, we reclassified the carbon AEFMs spanning less than 5-12 reactions depending on the GEM (shorter pathways), and the remaining carbon AEFMs (longer pathways). We found that longer pathways were associated with source-to-sink pathways, with shorter pathways containing the majority of looped pathways. In fact, 65-95% of all longer pathways in each GEM were source-to-sink pathways with metabolite revisitations, with this statistic increasing to 95-100% when including source-to-sink pathways without metabolite revisitations. Among shorter pathways, we observed that roughly half or more of them were looped carbon AEFMs with or without metabolite revisitations. Analysis of the nitrogen AEFMs revealed only a single distribution of relatively short pathways, since there are much fewer nitrogen-containing metabolites across all GEMs.

We next investigated how atoms within the same source metabolite may follow different paths through the network, irrespective of their AEFM class. We first computed the number of AEFMs for each carbon atom (resp. nitrogen atom). For each metabolite, we then computed the ratio of the class. We then computed the number of AEFMs for each carbon atom (resp. nitrogen atom). So for example, a ratio of 10 means that one of the carbon atoms in the metabolite can transit the network in 10 times as many different ways as the carbon with the fewest paths through the network. Our results are shown in Figure 3e-f, where each dot corresponds to the ratio for a given source metabolite. We found that a surprising number of carbons and nitrogens can transit different pathways within their respective source metabolites. Focusing on the source metabolites with carbon and nitrogen ratios over 100, we observed many of these corresponded to hydrophobic and positively-charged amino acids such as phenlylalanine, methionine, arginine, and histidine. Source metabolites containing fewer carbons or nitrogens tended to exhibit ratios closer to 1 with AEFMs involving the same metabolite subsets. Metabolites such as glycine, for example, are equivalently metabolized across the same carbon AEFMs. For each GEM, and not counting source metabolites with ratios of one, we computed median carbon ratios between 2.0 and 46.8 and nitrogen ratios between 1.3 and 11.2. Overall, these findings highlight the varying degrees of metabolic flexibility for atomic constituents of source metabolites to transit the metabolic network.

### A minority of carbon AEFMs explains the majority of source metabolite remodelling

We finally highlight the potential of our AEFM method for quantifying nutrient source metabolism. We turned to our HepG2 network which contained a single set of steady state fluxes estimated by Nilsson et al. These fluxes were estimated from a parsimonious FBA model of HepG2 cells cultured under exponential growth with external fluxes constrained by time-resolved exometabolomic measurements (see Methods for more information). Our focus was to analyze this flux network using our AEFM framework to identify alternative pathways of glutamine metabolism since many cancer cells, including HepG2, are dependent on this non-essential amino acid for survival and proliferation (Ye et al, 2023; Yoo et al, 2020). Using our method, we decomposed the set of steady state fluxes onto the 241,085 glutamine carbon AEFMs characterizing the HepG2 metabolic network. Figure 4a shows the cumulative explained mass flow of each glutamine carbon as a function of their AEFMs ranked from greatest to smallest explained mass flow. We capped the cumulative explained mass flow of each glutamine carbon at 99% because we observed that 98.9% of AEFMs collectively explained the remaining 1% glutamine carbon flow. These results suggested that the majority of glutamine carbon remodelling is explained by a surprisingly small number of glutamine AEFMs, especially since our Markovian flux decomposition approach assigns some weight to all network pathways. The top 5 ranked AEFMs across each glutamine carbon (25 pathways altogether) explained over half (50.2%) of the total glutamine carbon mass flow with the subsequent 5 next highest ranked AEFMs only explaining an additional 9.8% glutamine carbon mass flow. These observations were consistent across carbon AEFMs derived from other amino acids as well as glucose, leading us to hypothesize that steady state pathway fluxes are centralized to a relatively small subset of AEFMs (Figure S5). For some source metabolites, we observed overlapping curves between carbons (e.g. arginine, glycine, serine in Figure S5), suggesting that individual carbons were remodelled through the same AEFMs. However, corroborating results from Figure 3e-f, we observed many nonoverlapping curves for the majority of amino acids since their carbons could transit the network via different internal metabolites.

**Figure 4:**
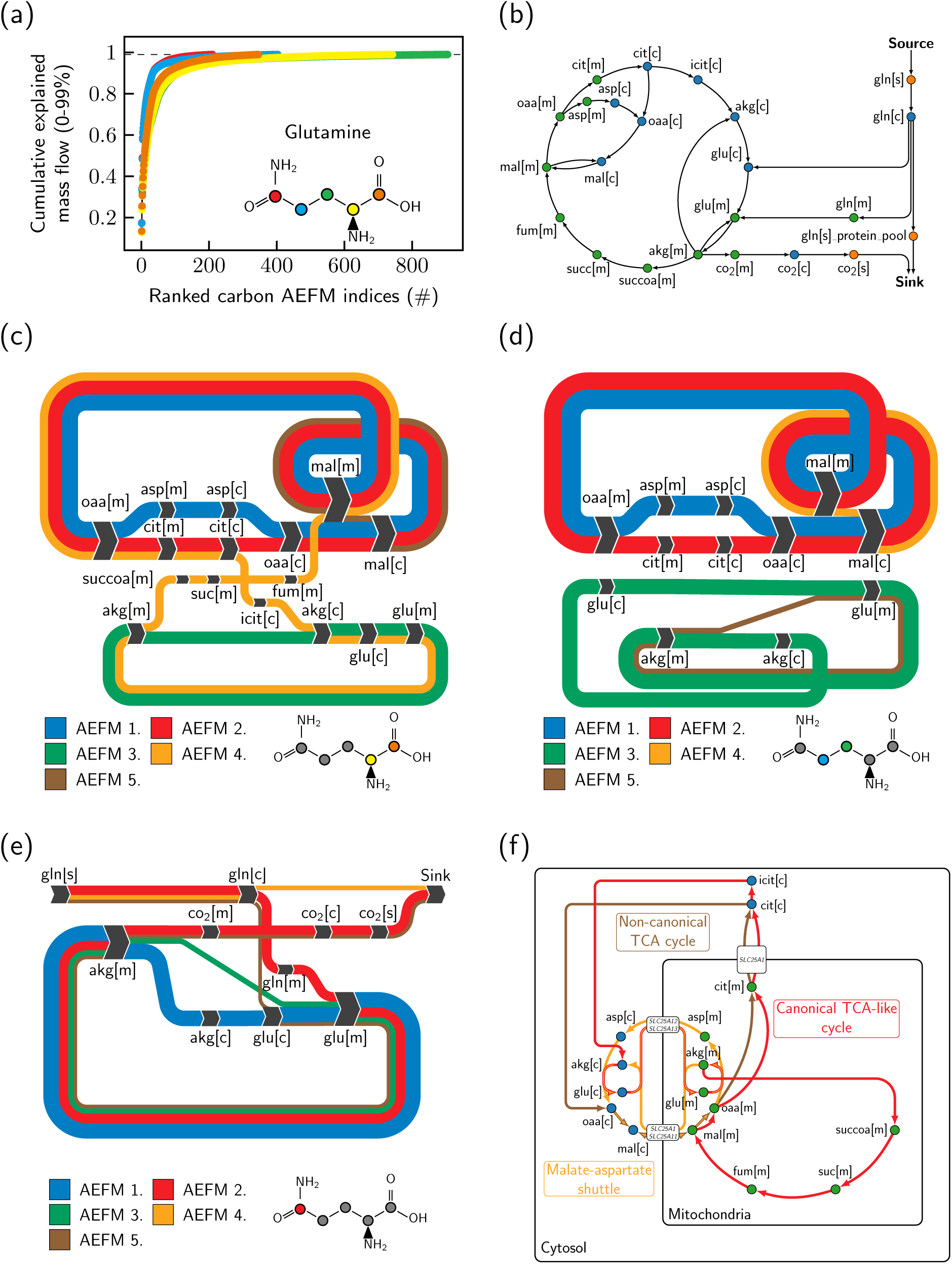
Carbon AEFM weight analysis of the HepG2 metabolic flux network. (a) A small number of highly active AEFMs explain the majority of glutamine flow. (b) Subnetwork of the top five AEFMs across each glutamine carbon collapsed onto metabolites. (c-e) Sankey diagrams of the top five glutamine carbon AEFMs. (f) Schematic mapping AEFMs onto traditionally defined metabolic pathways/subsystems. All metabolite names are labelled following the BiGG database nomenclature.

To obtain a high-level understanding of the dominant pathways of glutamine carbon remodelling, we constructed a metabolic subnetwork from the top 5 glutamine carbon AEFMs, collapsing metabolite-atom positions onto distinct metabolites, since none of these AEFMs involved metabolite revisitations. Of the 435 metabolites in the HepG2 dataset, input glutamine can remodel into 202 species localized to the cytosol or mitochondria. We found that the top 5 glutamine-derived carbon AEFMs consist of just 23 metabolites and 31 reactions (Figure 4b). These AEFMs generally consisted of cyclic pathways involving TCA intermediates. Only two source-to-sink reactions convert cytosolic glutamine towards the protein pool and carbon dioxide through decarboxylation of alpha-ketoglutarate in the mitochondria.

As AEFMs are steady state pathways, they can be visualized as Sankey diagrams to better understand the carbon flows through the network. Figure 4c-e shows the top 5 AEFMs for each glutamine carbon. Each AEFM in the diagram is shown as a flow proportional to its AEFM weight with nodes representing the metabolite transited by the given glutamine carbon. Although each glutamine carbon can transit a different number of AEFMs, we found the top 5 backbone carbons and first two sidechain carbons were identical in both AEFM pathway and weight, with no AEFM containing a metabolite revisitation.

Focusing on either backbone carbon in Figure 4c, we observed that all AEFMs corresponded to well-established metabolic subsystems. The first, third, and fifth highest ranked AEFMs corresponded to metabolites and reactions involved in the malate-aspartate shuttle. The fourth largest AEFM resembled a combination of canonical TCA cycle activity and malate-aspartate shuttle, with TCA intermediates citrate, isocitrate, and alpha-ketoglutarate localized to the cytosol rather than mitochondria. This AEFM also involved additional reactions transforming alpha-ketoglutarate into glutamate before remodelling into succinyl-CoA in the mitochondria. We noticed that the second largest AEFM cycled carbon mass between TCA intermediates citrate, oxaloacetate, and malate between the cytosol and mitochondria. Interestingly, this metabolic pathway corresponds to the recently discovered non-canonical TCA activity described by Arnold et al, who validated this pathway in non-small cell lung cancer, mouse embryonic, and C2C12 mouse myoblast cells by stable isotope tracing analysis (Arnold et al, 2022). This non-canonical TCA activity was further observed in glutamine carbons 3 and 4 in AEFM 2 (Figure 4d). We also found AEFMs 1 and 3 in Figure 4d-e corresponded to the malate-asparate shuttle. Only the terminal glutamine carbon in Figure 4e contained source-to-sink AEFMs that decarboxylated glutamine-derived alpha-ketoglutarate into carbon dioxide.

Our analyses revealed that the most active glutamine-derived carbon AEFM weights corresponded to well-known metabolic subsystems. This finding led us to ask whether we could summarize our AEFM analysis as a simplified metabolic model describing the majority of glutamine carbon remodelling in the HepG2 flux network. The resulting schematic is shown in Figure 4f which consists of the following three major metabolic pathways: canonical TCA-like cycle, non-canonical TCA cycle, and the malate-aspartate shuttle. These metabolic pathways were chosen based on the number of top glutamine-derived carbon AEFMs that overlapped with reactions within these subsystems. Our results suggest input glutamine is converted into cytosolic glutamate which is transported into the mitochondria through aspartate/glutamate antiporters encoded by *SLC25A12* and *SLC25A13* annotated in the HepG2 metabolic model. Mitochondrial glutamate is then converted into TCA intermediate alpha-ketoglutarate and citrate. Instead of following the canonical reactions of the TCA cycle, the mitochondrial citrate is exported to the cytosol where it can participate in noncanonical TCA cycle activity or the malate-aspartate shuttle to re-enter the mitochondria. Taken together, the dominant carbon AEFMs predicted by our model suggest HepG2 glutamine is conserved within TCA intermediates and regenerated through subsequent reactions involving cytosolic citrate.

## Discussion

Since the proposal of molecular EFMs, much effort has been made to enumerate and decompose steady state metabolic fluxes onto these pathways. By defining EFMs with respect to individual atoms rather than molecules, we introduced the concept of AEFMs which describe how individual atoms are remodelled within a metabolic network. As AEFMs never involve branching pathways, they are easier to interpret than their molecular EFM counterparts, which may involve multiple inputs and outputs to balance molecular stoichiometries. Using the atom mapping program RXNMapper, we adapted our previously described CHMC method to efficiently enumerate carbon and nitrogen AEFMs in five GEMs. Our structural analysis systematically characterized source metabolite remodelling at the atomic level. By comparing the number of AEFMs across atoms within the same source metabolite, we revealed how certain amino acids displayed remarkable variability in the number of pathways transited by their constituent atoms. This analysis, in particular, may be useful for designing non-uniformly labelled stable isotope tracers to identify fluxes within specific metabolic subsystems. Our AEFM weight analysis of the HepG2 network revealed that the majority of glutamine carbons were remodelled via cyclic pathways regenerating TCA intermediates through cytosolic citrate. Excitingly, our model also predicted non-canonical TCA activity which has not been previously described in HepG2 cells.

As a systematic method to explain the flow of metabolites, the problem of molecular EFM enumeration has historically limited their applicability to small-scale networks. Many algorithmic advances have incrementally improved molecular EFM enumeration, yet never feasibly beyond networks with *>* 100 metabolites and reactions (Kamp & Schuster, 2006; Terzer & Stelling, 2008; van Klinken & Willems van Dijk, 2016). Molecular EFM enumeration is generally regarded as infeasible for large-scale networks—so much so that numerous methods have been developed to enumerate a subset of molecular EFMs constrained by pathway length (de Figueiredo et al, 2009), reaction thermodynamics (Gerstl et al, 2015), participating metabolites (Pey & Planes, 2014), participating reactions (Kaleta et al, 2009), gene regulatory rules (Jungreuthmayer et al, 2013), metabolic subsystems (Schuster et al, 2002; Schwartz et al, 2007), extracellular flux measurements (Soons et al, 2011; Jungers et al, 2011), or even random sampling (Machado et al, 2012). While our HepG2 results did show the majority of source metabolite fluxes were explained by a small number of AEFMs, contextualizing the significance of these dominant pathway fluxes is only possible when all AEFMs are known. Furthermore, reaction thermodynamics, gene regulatory rules, and other constraints may change across different biological conditions. By computing AEFMs, instead, we could feasibly enumerate atomic pathways in genome-scale networks containing hundreds of metabolites and reactions within minutes or hours on consumer computing hardware. We are hopeful that algorithmic tricks borrowed from molecular EFM enumeration tools, such as linear pathway compression, could further decrease memory usage and run times.

Aside from carbon and nitrogen AEFMs, our work is broadly applicable to other types of chemical elements. One could feasibly enumerate oxygen or hydrogen EFMs in small-scale networks, although the significance of these pathways is unclear for biochemical networks given the ubiquitous nature of these atoms. We also caution against enumerating hydrogen AEFMs, in particular, given limitations associated with RXNMapper. Unless explicitly represented in the reaction SMILES string, RXNMapper does not assign hydrogen atom mappings. Moreover, RXNMapper cannot map atoms in reaction SMILES strings longer than 512 characters. Our method could also be applied to study sulphur metabolism to quantify methionine or cysteine remodelling, or phosphorus metabolism to quantify nucleotide synthesis. Outside the realm of biochemistry, our method is applicable to any periodic table element and possibly relevant for studying general chemical reaction networks.

Our AEFM framework opens many new computational problems in metabolism. While we analyzed one set of fluxes for the HepG2 network, one could envision a differential AEFM analysis of flux distributions under differential conditions. These differential AEFMs may reveal changes in metabolic subsystem activity or predict new pathways of source metabolite remodelling. For instance, analyzing carbon AEFMs of C1 compounds may be particularly useful for quantifying or optimizing carbon fixation pathways in C1-utilizing organisms. Another important question is whether a joint analysis of AEFM weights could reveal broader metabolic changes within a condition. We observed that many AEFMs across distinct source metabolites converged on common internal metabolites. Intuitively, the weights of AEFMs rooted on different source metabolites, but sharing common downstream metabolite subsequences, should be related, and these relationship may further speed AEFM enumeration or weight computations. Finally, more work is needed to establish relationships between AEFM decompositions and more traditional EFM decompositions of fluxes, and to explore the possibility that feasible AEFM calculations might somehow aid in EFM calculations.

## Materials and Methods

### Methods and Protocols

#### Datasets

Five highly-curated GEMs were chosen in this study to reflect a diversity of organisms, metabolites and biochemical reactions. Four networks were obtained from the BiGG UCSD database (King et al, 2016) in ascending order of network size. Datasets were downloaded as SBML (.xml) files, and the corresponding stoichiometry matrices, metabolites and reactions were extracted using the Julia SBML.jl package. The HepG2 liver cancer cell line dataset was chosen for its single set of estimated steady state fluxes estimated from a FBA with external fluxes constrained by experimental measurements of amino acid, glucose, pyruvate, and lactate. These cells were grown in DMEM with 10% fetal calf serum, 22mM glucose, and 1.8mM glutamine (Nilsson et al, 2020b). The steady state fluxes were computed from the FBA model provided by the authors (figure3 metabolitebalances.m; MATLAB R2018a) (Nilsson et al 2020b; Data ref: Nilsson et al 2020a). Across all models, we modelled all reversible reactions as net forward reactions to simplify the number of molecular and AEFMs as the majority of experimental and computational methods can only estimate net fluxes. We assumed the reaction stoichiometries of the BiGG datasets represented the canonical net directions of all reversible reaction. For the HepG2 dataset, the net directions were chosen based on the net fluxes estimated from the FBA model.

#### Extracting metabolites SMILES strings

SMILES structures were not encoded in the BiGG SBML files nor the HepG2 dataset. Thus, we manually extracted SMILES strings corresponding to each metabolite across all GEMs. Isomeric SMILES strings were taken from PubChem (Kim et al, 2023), the Human Metabolome Database (Wishart et al, 2022), or MetaNetX database (Moretti et al, 2021). Metabolites shared across GEMs were assigned the same SMILES structure in addition to identical metabolites stratified across distinct subcellular compartments. Generic structures such as R-groups were not modelled in this study and were replaced with a single bonded hydrogen under the assumption this modification would not alter atom mappings, especially since RXNMapper does not map hydrogen atoms unless they are explicitly represented in the SMILES strings. SMILES strings were also not assigned to metabolites with ambiguous structures nor pseudometabolites with no known chemical structure. All pseudometabolites and reactions involving pseudometabolites were removed from the stoichiometry matrix (Table 2).

**Table 2:** Number of metabolites and reactions removed from each network during preprocessing.

|  | E. coli core | iAB_RBC_283 | iIT341 | iSB619 | HepG2 |
| --- | --- | --- | --- | --- | --- |
| Rxns with non-integer stoichiometries | 2 | 1 | 3 | 34 | 8 |
| Pseudometabolites with no known structures | 1 | 68 | 52 | 108 | 33 |
| Rxns with pseudometabolites | 2 | 142 | 93 | 204 | 46 |
| Rxns exceeding RXNMapper token limit | 0 | 1 | 4 | 14 | 3 |

#### Constructing reaction SMILES strings

RXNMapper performs atom mapping on a string that encodes the SMILES structures of all substrates and products for a given reaction. These reaction SMILES strings are constructed by concatenating SMILES strings of individual substrates and products. Multiple substrates or products are separated by the symbol ‘.’, while the reaction arrow delimiting substrates and products is denoted by the symbol ‘*>>*’. The substrate and product SMILES strings are concatenated in order of metabolite appearance within the stoichiometry matrix, and multiple stoichiometric copies are consecutively repeated in the reaction SMILES strings. Non-integer stoichiometric coefficients are therefore not allowed and these pseudoreactions were removed from the stoichiometry matrix (see Table 2). To maintain steady state flux in the HepG2 network, flux along pseudoreactions were replaced with unimolecular source and sink fluxes consuming each pseudoreaction substrate and producing each pseudoreaction product. We manually checked several reactions to ensure the assigned atom mappings were correct. Crucially, we found RXNMapper to fail on nearly all classes of ATP-independent transaminase reactions. This problem has previously been reported in an atom mapping benchmark study prior to the publication of RXNMapper (Preciat Gonzalez et al, 2017). These transaminase reactions were identified in all metabolic networks and manually atom mapped based on the KEGG reaction mappings (Figure S6).

#### Pre-processing genome-scale metabolic models

Several pre-processing steps were required to compute AEFMs from the GEMs. The stoichiometric coefficients of external reactions were set to one and their fluxes rescaled to carry equivalent units of source and sink metabolites. All reactions containing pseudometabolites with no known SMILES strings were removed from the stoichiometry matrix and replaced with unimolecular source and sink reactions carrying equivalent units of flux. The same procedure was also applied to reactions containing non-integer stoichiometries. These pseudometabolites were next removed from the stoichiometry matrix rows, and unimolecular reactions involving the same external or internal metabolites were aggregated. Finally, we had to remove a small number of reactions whose reaction SMILES strings exceeded the 512 token limit of RXNMapper. These reactions were likewise converted into unimolecular source and sink reactions to maintain steady state flux across the network (see Table 2). All pre-processing steps described above are wrapped into convenience functions within our Julia package MarkovWeightedEFMs.jl.

#### Benchmarking enumeration of atomic-versus-standard EFM

We benchmarked our Julia package MarkovWeightedEFMs.jl version 2.0 running Julia version 1.10.2 to enumerate AEFMs versus the molecular EFM enumeration program FluxMod-eCalculator (van Klinken & Willems van Dijk (2016)) running MATLAB version R2018a with default parameters including network compression. We chose FluxModeCalculator because it is the state-of-the-art program for enumerating molecular EFMs. Benchmarks were performed on a 64-bit Linux operating system with 64GB of memory and an AMD 5950X CPU with 16 cores/32 threads. All benchmarks reported the wall time to run the atomic or molecular EFM enumeration functions and did not include startup times to open Julia/MATLAB, precompile Julia functions, or load packages and input data. A benchmark time of ‘did not finish’ (DNF) was assigned if either program could not complete (A)EFM enumeration within 7 days. In practice, only FluxModeCalculator failed to enumerate the molecular EFMs in three of the five datasets.

#### Constructing atomic transition graphs

For each carbon and nitrogen atom within a given source metabolite, we identify all possible metabolite/atom states that are traversable by that initial state and constrained by the reaction atom mapping predictions. For reactions involving single stoichiometric copies of each substrate and product, the atom of interest will always move from a single substrate to product molecule. If this atom is present in a substrate with multiple stoichiometric copies, we assume there is an equal probability of that atom occurring in either copy. Depending on the substrate stoichiometric copy, that atom may move to (i) different product metabolite/atom states, (ii) the same product but at different atomic positions, or (iii) the same metabolite/atom position. In all cases, we set the atomic flux of the atom transitioning from either stoichiometric copy equal to the reaction fluxes to maintain atomic mass conservation (Figure S1). The resulting atomic transition graph is completed by connecting any sink fluxes to the root of the tree initialized on the source metabolite/atom position.

#### ACHMC model

Following Chitpin & Perkins, the atomic transition graphs were analyzed by an atomic version of our CHMC (ACHMC) method to enumerate the corresponding AEFMs and compute their weights. In this formulation, each ACHMC state corresponds to a sequence of metabolite-atom positions transited by a given source metabolite atom. Each simple cycled identified by the ACHMC corresponds to an AEFM rooted on a user-specified source metabolite atom. For the HepG2 flux network, we computed the probabilities of each AEFM by steady state analysis of the ACHMC. These AEFM probabilities were scaled to weights such that the totality of source-to-sink AEFMs was equal to the unimolecular input flux producing that source metabolite.

## Data availability

Our software package MarkovWeightedEFMs.jl is available at https://github.com/jchitpin/ MarkovWeightedEFMs.jl. All source codes to reproduce this study are available at https://www.github.com/jchitpin/reproduce-efm-paper-2024.

## Acknowledgements

This research was supported by the Natural Sciences and Engineering Research Council of Canada (NSERC), Discovery grant RGPIN/06604-2019 and a Digital Research Alliance of Canada (DRAC) Resources for Research Groups grant to TJP (RRG-TPERKINS). J.G.C. was supported by an NSERC CREATE Matrix Metabolomics Scholarship, NSERC Alexander Graham Bell Canada Graduate Scholarship, and Ontario Graduate Scholarship.

## Author contributions

Justin G. Chitpin: Conceptualization; data curation; methodology; software; formal analysis; investigation; visualization; writing - original draft; writing - review and editing.

Theodore J. Perkins: Conceptualization; methodology; supervision; funding acquisition; investigation; project administration; writing - review and editing.

---

**Figure S1:**
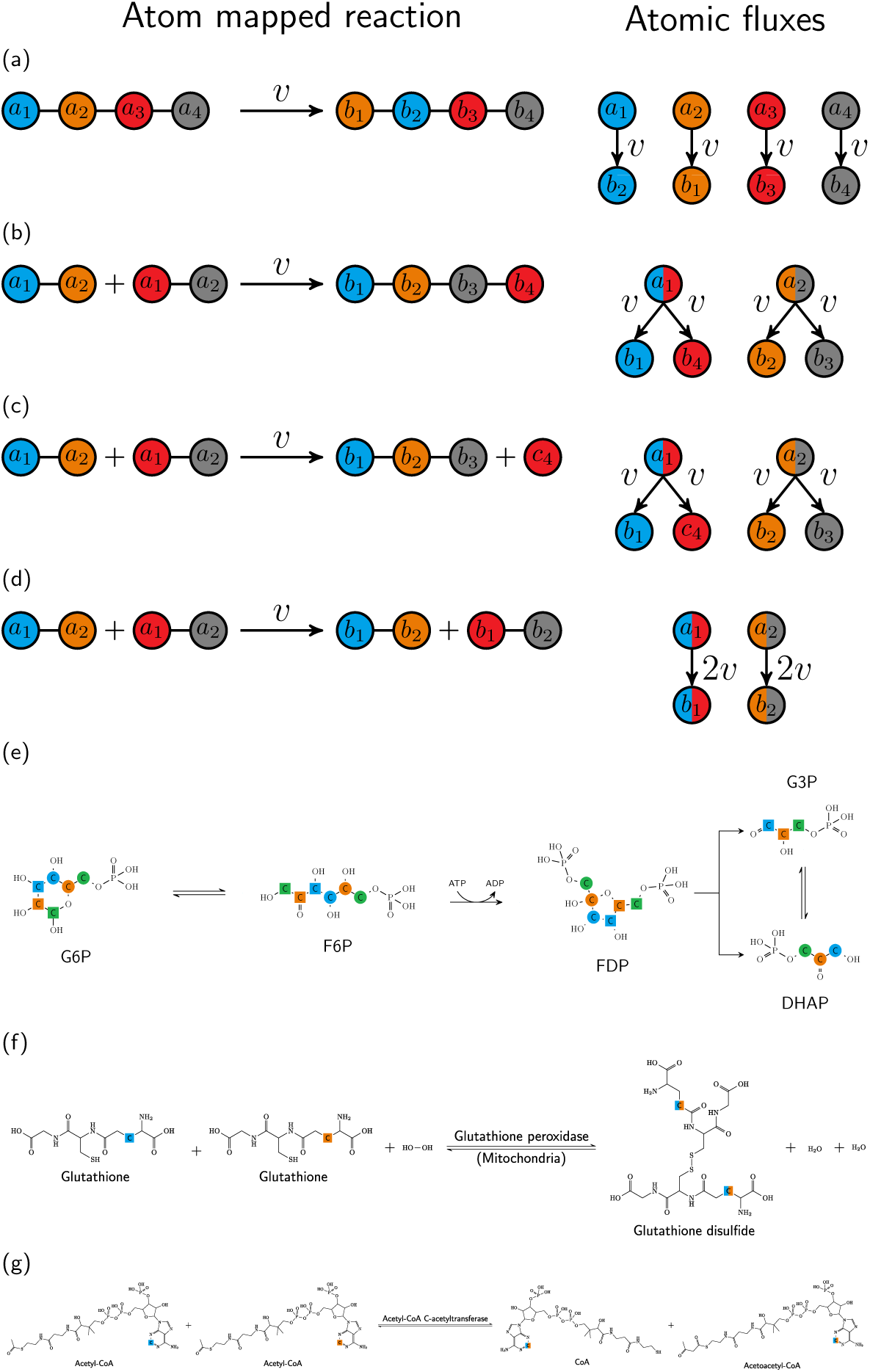
Rules for assigning atomic fluxes from atom-mapped reactions with biological examples. Schematic node colours represent ground truth atom mappings across substrate(s) and product(s). Node labels correspond to distinct atomic indices of substrate and product stoichiometric copies. (a) For reactions with single stoichiometric copies of substrates and products, each atom within a given substrate moves to a single atomic position within a product with atomic flux equal to the reaction flux. (b-d) For reactions with *s* stoichiometric copies of a given substrate, there are *s* copies of each metabolite-atom state on the substrate and product side of the reaction. The probability of any atom within a substrate stoichiometric copy moving to a product metabolite-atom state remains equal to the reaction flux. (b) The stoichiometric copies of the substrate fuse together into a single product. (c) The stoichiometric copies of the substrate form distinct products. (d) The *s* stoichiometric copies of the substrate form *s* copies of products with identical atomic backbones. (e) Biological example of rule a where the atoms in each substrate map to a single position on the corresponding product. (f) Biological example of rule b where two stoichiometric copies of glutathione fuse together to form a single stoichiometric copy of glutathione disulfide. (g) Biological example of rule c where two stoichiometric copies of acetyl-CoA undergo a reaction to produce a stoichiometric copy of two distinct products.

**Figure S2:**
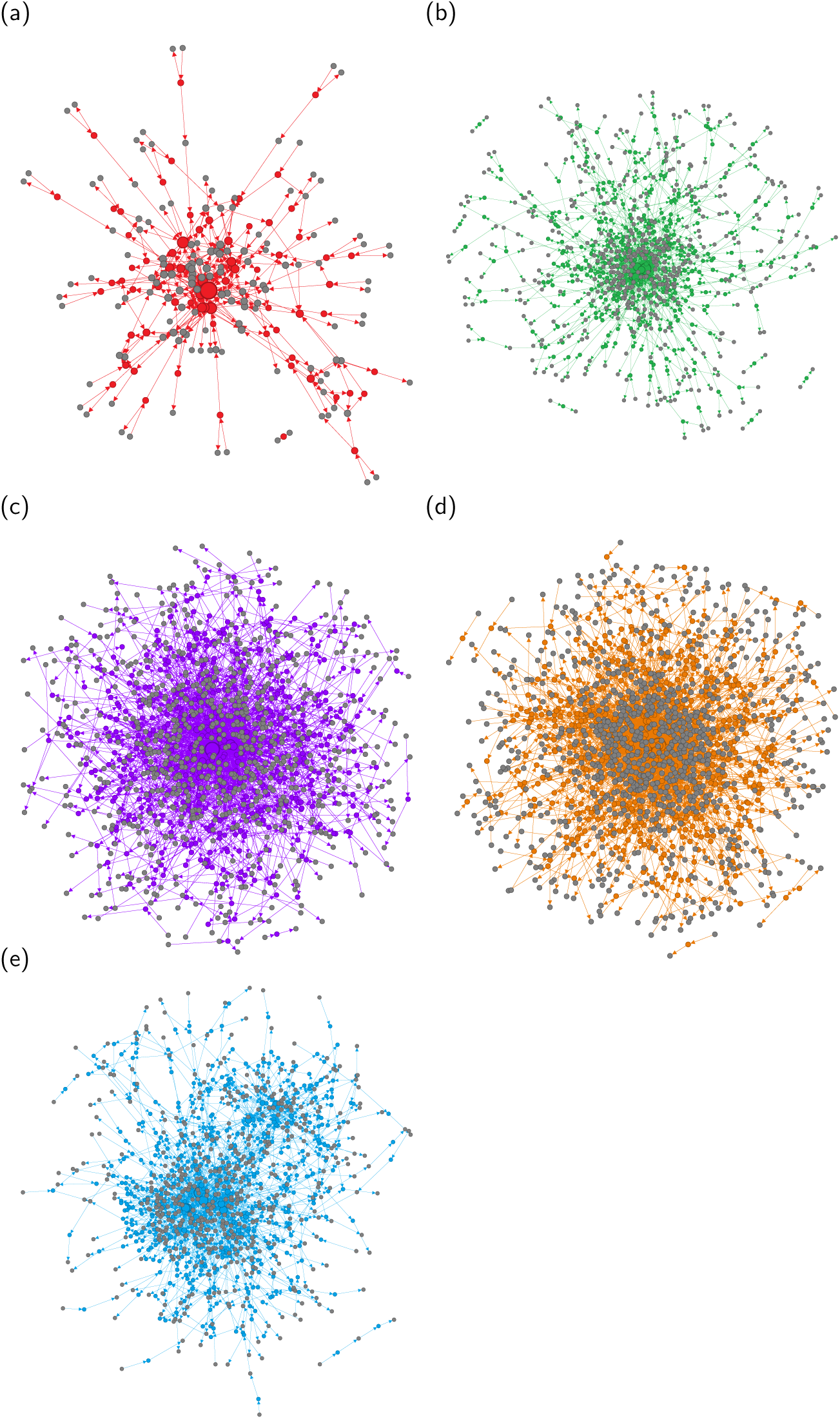
Network visualization of the five GEMs after pre-processing. Metabolites (non-grey nodes) are connected to reaction entities (grey nodes) with node sizes proportional to their degree. Graphs were constructed in Gephi version 0.10 using the Yifan-Hu proportional network layout with default parameters. (a) E. coli core. (b) iAB RBC 283. (c) iIT341. (d) iSB619. (e) HepG2.

**Figure S3:**
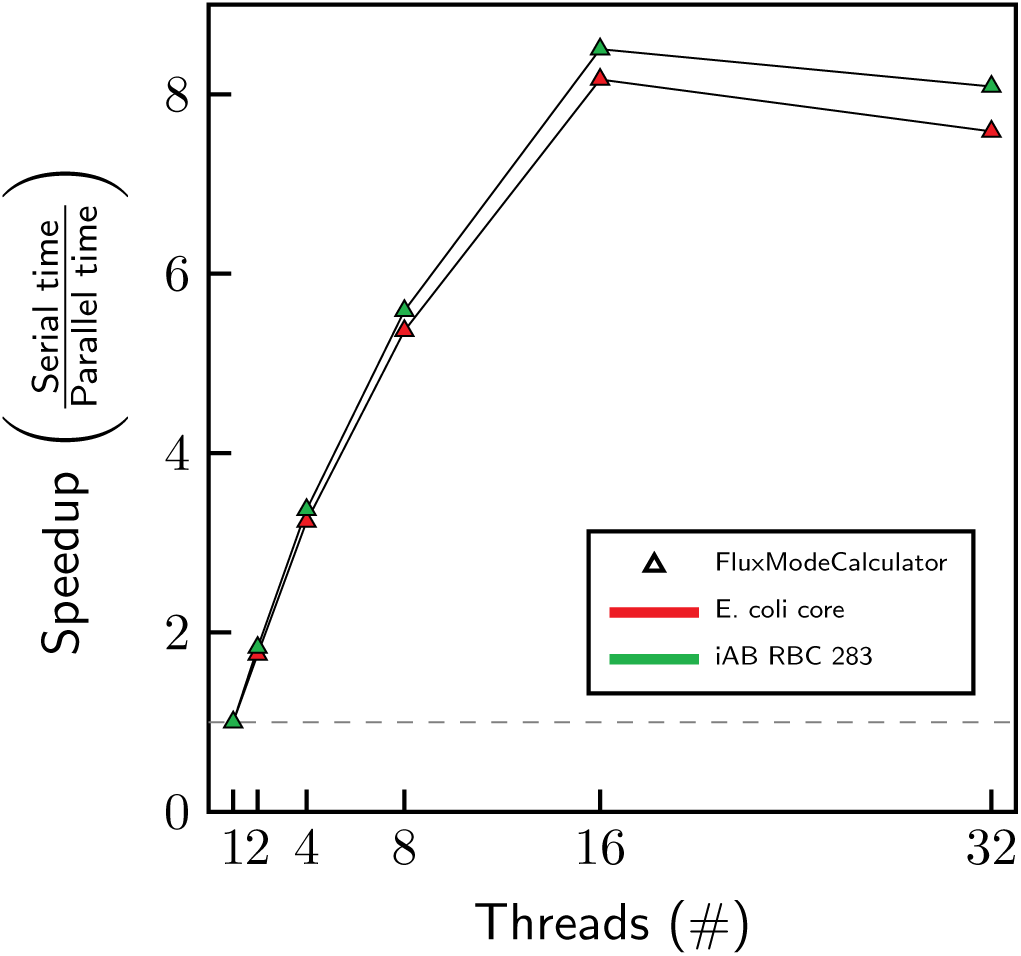
FluxModeCalculator scaling across multiple threads.

**Figure S4:**
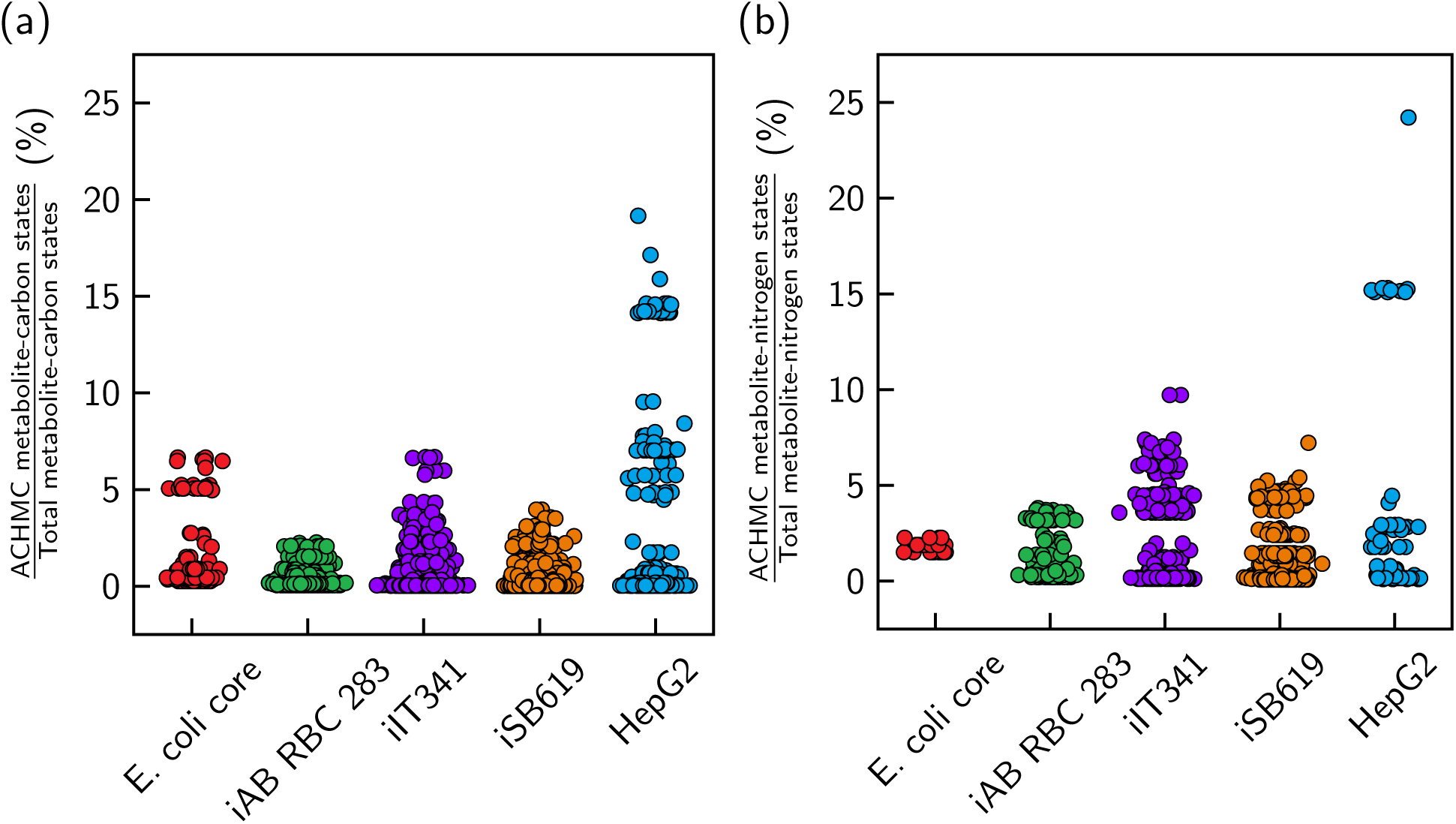
Most ACHMCs traverse less than 10% of the total metabolite-carbon/nitrogen state space. Each point corresponds to an ACHMC rooted on a given source metabolite carbon or nitrogen.

**Figure S5:**
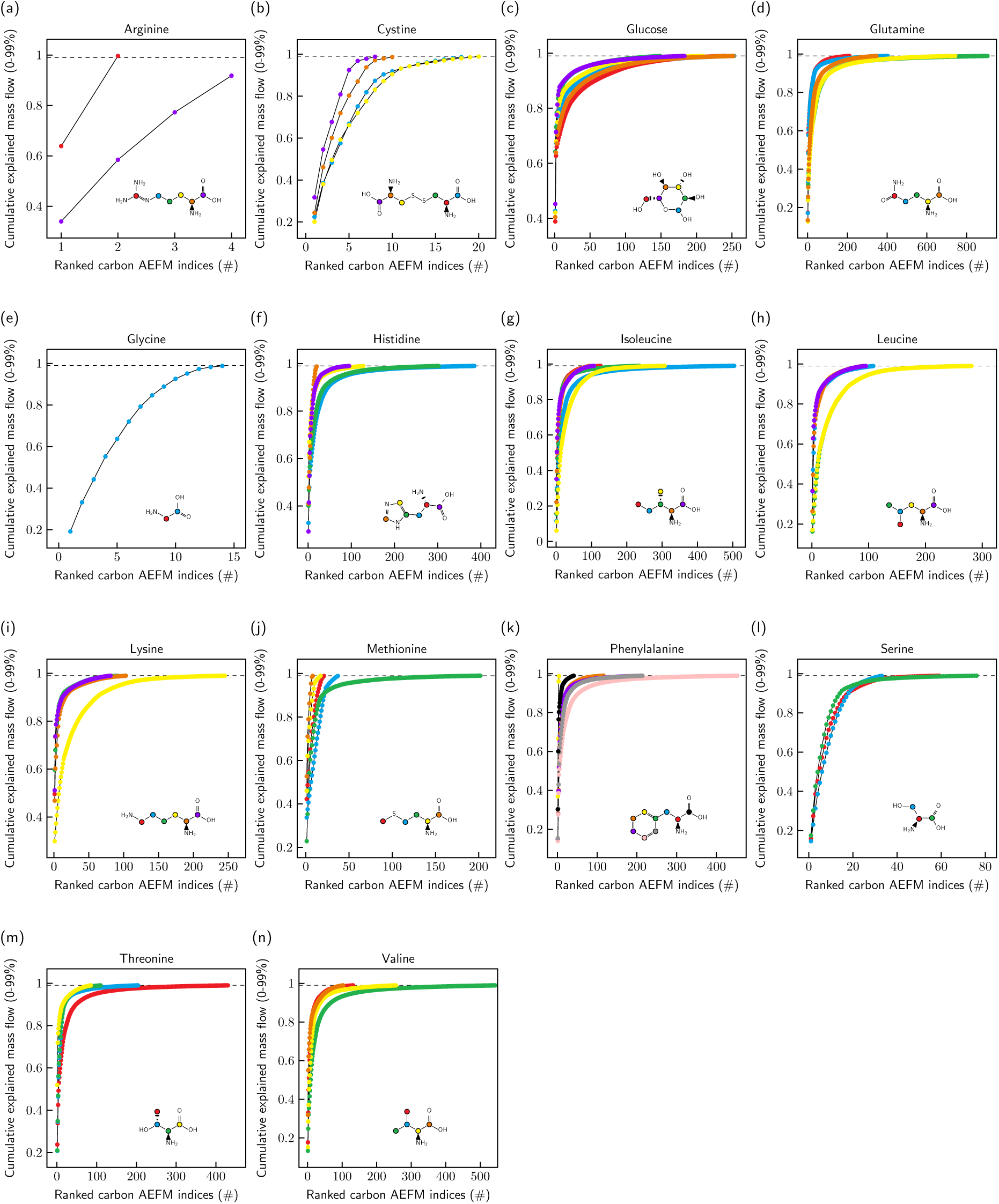
Cumulative explained carbon mass flow of input glucose and amino acid carbons from the HepG2 flux network. The cumulativeexplained mass flow is truncated at 99% to avoid displaying AEFMs that carry little atomic mass flow.

**Figure S6:**
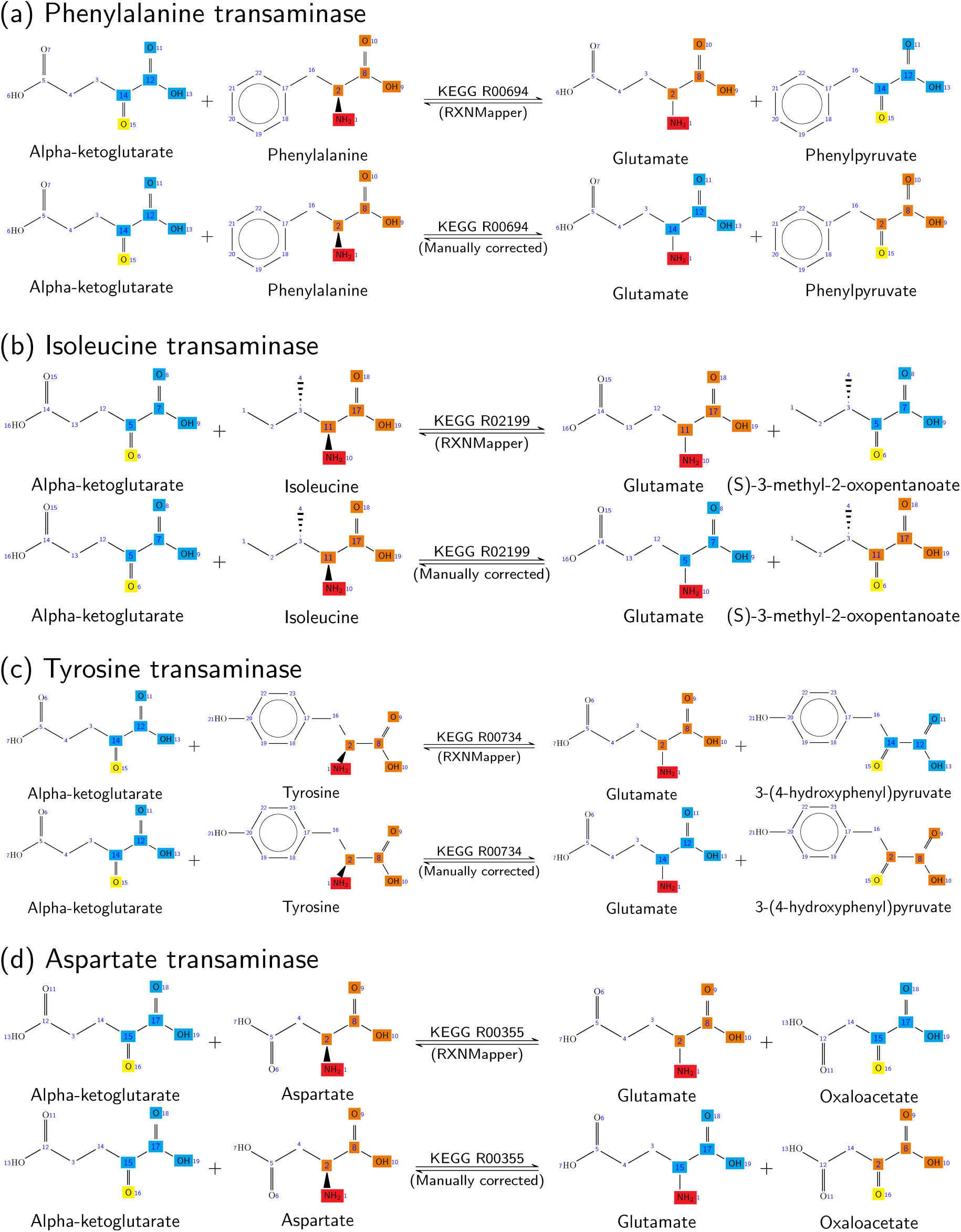
Examples of incorrectly atom-mapped transaminase reactions present in one or more of the five networks. (a) Phenylalanine transaminase. (b) Isoleucine transaminase. (c) Tyrosine transaminase. (d) Aspartate transaminase.

